# *In vivo* imaging of coral tissue and skeleton with optical coherence tomography

**DOI:** 10.1101/088682

**Authors:** Daniel Wangpraseurt, Camilla Wentzel, Steven L. Jacques, Michael Wagner, Michael Kuhl

## Abstract

Optical coherence tomography (OCT) is a non-invasive three-dimensional imaging technique with micrometer resolution allowing microstructural characterization of tissues *in vivo* and in real time. We present the first application of OCT for *in vivo* imaging of tissue and skeleton structure of intact living corals spanning a variety of morphologies and tissue thickness. OCT visualized different coral tissue layers (e.g. endoderm vs ectoderm), special structures such as mesenterial filaments and skeletal cavities, as well as mucus release from living corals. We also developed a new approach for non-invasive imaging and quantification of chromatophores containing green fluorescent protein (GFP)-like host pigment granules in coral tissue. The chromatophore system is hyper-reflective and can thus be imaged with good optical contrast in OCT, enabling quantification of chromatophore size, distribution and abundance. Because of its rapid imaging speed, OCT can also be used to quantify coral tissue movement showing that maximal linear contraction velocity was ~120 μm per second upon high light stimulation. Based on OCT imaging of tissue expansion and contraction, we made first estimates of dynamic changes in the coral tissue surface area, which varied by a factor of 2 between the contracted and expanded state of the coral *Pocillopora damicornis*. We conclude that OCT is an excellent novel tool for *in vivo* tomographic imaging of corals that can reveal tissue and skeleton organization as well as quantify dynamic changes in tissue structure and coral surface area non-invasively and at high spatio-temporal resolution.

## Introduction

Coral reefs are hotspots of biodiversity and one of the most productive ecosystems on Earth (Hatcher, 1990; Odum and Odum, 1955). The key drivers of this productive ecosystem are reef building (scleractinic) corals, i. e., invertebrates that belong to the family Cnidaria, that live in symbiotic interaction with dinoflagellate microalgae of the genus *Symbiodinium*. Coral reefs are characterized by a rich structural complexity as coral colonies grow in diverse shapes such as branch-like, hemispherical, encrusting or plate-like morphotypes (Kaandorp and Kübler, 2001). The structure of coral skeletons has been studied in detail from macroscopic colony scales down to the nanometer scale, where scanning electron microscopy, 3D laser scanning and x-ray microcomputed tomography have acquired high resolution structural information of coral skeletons (Meibom et al., 2008; Raz-Bahat et al., 2009; Stolarski, 2003), which has significantly advanced our understanding of coral structure and calcification (Tambutté et al., 2007). Coral skeleton microstructure has also been shown to modulate the backscattering of light (Marcelino et al., 2013), which can affect the light exposure of *Symbiodinium* in the coral tissue (Enriquez et al., 2005), and it has been suggested that such skeleton-derived differences in backscatter can explain differences among corals in their susceptibility to bleaching under environmental stress (Swain et al., 2016).

Much less is known about the structural complexity of intact living corals, albeit coral tissue and its plasticity evidently must play an important role for coral ecophysiology. The exchange of solutes and metabolites is controlled at the coral tissue-water interface and is a function of the interaction between water flow and coral tissue topography (Kühl et al., 1995; Wangpraseurt et al., 2012b). Coral tissue microtopgraphy affects the thickness of the diffusive boundary layer (DBL), that is the thin layer of water covering the coral tissues where diffusion is the prime transport mechanism, which can be rate limiting for respiration and photosynthesis (Jørgensen and Revsbech, 1985; De Beer and Kühl, 2001; Larkum et al., 2003). The buildup of a thermal boundary layer (TBL) controls the heat exchange of corals and is likewise affected by flow and tissue surface topography (Jimenez et al., 2011). Additionally, coral tissue properties strongly modulate the optical microenvironment of coral zooxanthellae, where e.g. tissue thickness and opacity control *Symbiodinium* light exposure via formation of distinct light gradients (Wangpraseurt et al., 2012a) that are further affected by the presence and distribution of coral host pigments (Lyndby et al., 2016; Salih et al., 2000). The thermal microenvironment of *Symbiodinium in hospite* is also affected by tissue optical properties (Jimenez et al., 2012; Lyndby et al., 2016).

The structural properties of coral tissues are not static, and corals can re-arrange their tissues in response to the ambient light environment (Levy et al., 2003; Wangpraseurt et al., 2014), changes in the concentration of gases (Kühl et al., 1995), and water movement and food exposure (Sebens and Johnson, 1991). Both the thermal and optical microenvironment of *Symbiodinium* are affected by tissue contraction or expansion (Lyndby et al., 2016; Wangpraseurt et al., 2014). Together, the structural and optical properties of the coral tissue and its plasticity in terms of distribution over the coral skeleton strongly modulate coral photosynthesis, respiration, and the exchange of solutes with the surrounding seawater (Patterson 1992), and there is a need for non-invasive *in vivo* imaging techniques that can resolve tissue structural dynamics at high spatio-temporal resolution and sufficient areal coverage.

Descriptions of coral tissue structure have largely been based on light microscopy (Brown et al., 1995; Hayes and Bush, 1990), histological sectioning, and electron microscopy, which requires extensive tissue sample preparation that can introduce artefacts such as tissue shrinkage and dehydration. Microscopic imaging of live intact corals has also been realized in the lab (e.g., Shapiro et al., 2016; Sivaguru et al., 2014) and *in situ* (Mullen et al., 2016) but information about internal tissue organization is limited to early developmental stages, tissue explants or corals with thin tissue layers without pronounced calcified structures. Depth resolution and contrast is severely hampered by the absorption and scattering of visible light in the tissue-skeleton matrix, and the same effects contrive the depth resolution of confocal laser microscopy on coral tissue to the outermost 100-200 μm using high numerical aperture objectives with a small field of view (Salih, 2012). Recently, digital holographic microscopy (DHM) was used for non-invasive imaging of mucus formation in cold-water corals (Zetsche et al., 2016). DHM is a novel, essentially lense-free phase imaging method based on recording the interference between visible light passing through the sample and a reference light beam, where the interference pattern of an imaged object is first stored as a digital hologram and then digitally processed with reconstruction algorithms (Zetsche et al., 2014 and references therein). While DHM can provide fast and non-invasive imaging of coral surfaces and (semi-)transparent surface layers such as mucus with a larger depth of focus as compared to normal light microscopy, its application for deeper tomographic imaging of opaque coral tissue and skeleton structures is limited.

Biomedical tissue imaging faces similar challenges as listed above for coral tissues and several spectroscopic imaging techniques have been developed, where the use of near infrared radiation (NIR) bears distinct advantages for non-invasive structural imaging in terms of penetration depth, areal coverage and image acquisition speed (Vogler et al., 2015). A key technique that has revolutionized e.g. dermatology and ophthalmology is optical coherence tomography (OCT) (Huang et al., 1991), that allows for non-invasive *in vivo* cross sectional characterization of tissues with micrometer resolution using NIR. OCT is often referred to as an optical analogue to ultrasound, which measures the echo time delay of sound waves travelling into materials, where they are backscattered according to certain frequencies that depend on the acoustic properties of different material components. Likewise, OCT measures characteristic patterns of directly elastically backscattered (low coherent ballistic and near ballistic) photons from different reflective layers in a sample, e.g., at refractive index mismatches between tissue compartments with different microstructural properties (Schmitt, 1999). In the present study, we explored the use of OCT for studying coral tissue microstructure and dynamics. We demonstrate that OCT can be used as a rapid, non-invasive coral imaging technique that can easily be implemented in experimental setups to gain high resolution tomographic data on coral skeleton and tissue structure, and we present novel OCT-based approaches for the *in vivo* characterization of a green fluorescent protein (GFP)-like chromatophore system, tissue movement and surface area in living corals.

## 2. Methods

### 2.1. OCT system and measuring principle

We used a commercially available spectral-domain (SD) OCT system (Ganymede II, Thorlabs GmbH, Luebeck, Germany) equipped with an objective lens with an effective focal length of 36 mm, and a working distance of 25.1 mm (LSM03; Thorlabs GmBh, Luebeck, Germany; Fig. 1a). The OCT system includes a superluminescent diode (SLD) emitting broadband low coherent light centered at 930 nm. The light source sets a limit to the axial resolution of OCT imaging defined as the so-called coherence length or gate (Fercher et al., 2003):

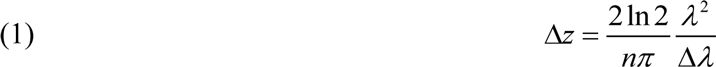

where *n*= the refractive index of the medium (*n* = 1.33 for water), *λ* is the center wavelength (*λ* = 930nm) and *Δλ*is the full width at half maximum of the power spectrum of the SLD (*Δλ*=65nm). Thus in our case *Δz* was 4.5 μm. In contrast, the lateral resolution is dependent on the imaging optics, which generated a limit to the *x-y* resolution of 8 μm.

**Fig. 1.**
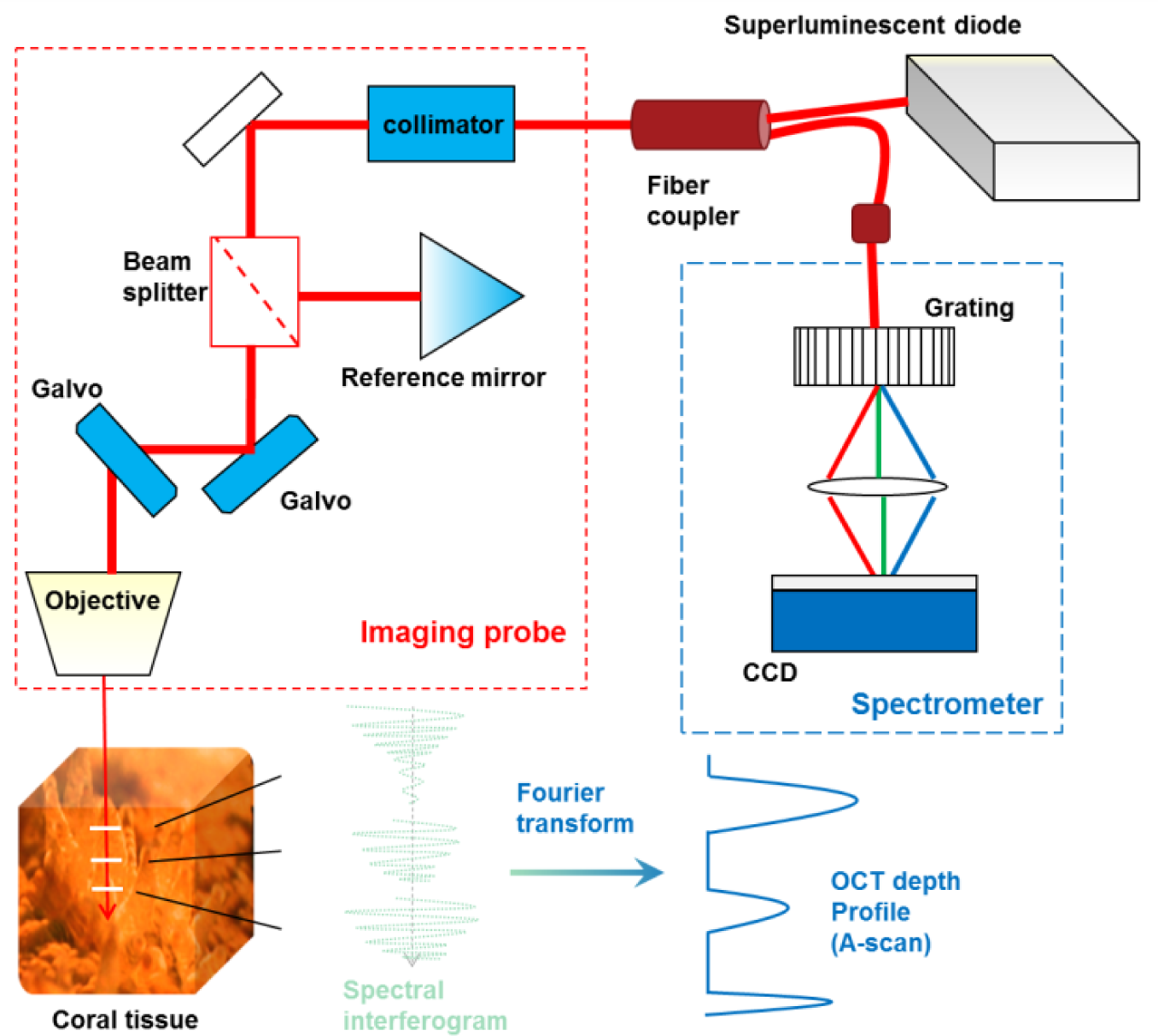
Principle of spectral-domain optical coherence tomography. The super-luminescent diode provides broadband low coherent near infrared radiation (930 nm). Light is emitted through the fiber and send to the imaging probe, which works like a modified Michaelson interferometer. The beam splitter sends light to the retro-reflecting prism (reference mirror) and the sample beam which is focused by the scanning objective before it interacts with the coral tissue. Reflective boundaries within the coral tissue (white lines) scatter light back to the imaging probe where, reference light and sample light are combined and send to the spectrometer. A diffraction creating creates a characteristic spectral interferogram (Fourier-domain signal), which is converted to an OCT depth profile of reflectivity along the z-axis (A-scan) via Fourier transform. Two-dimensional (OCT B-scan) and three-dimensional (OCT C-scan) tomographs are sampled by moving the sample beam within the imaging probe along the *x* and *y* axis by a galvanometer system.

The light generated by the SLD is coupled into a bifurcated fiber which is connected to the OCT imaging probe (Fig. 1). The key components of the imaging probe consist of a beam splitter, a galvo system (galvanometer and mirror), a retro-reflecting prism and a charged coupled device (CCD) detector (Fig. 1). The OCT system is additionally equipped with a USB camera and LED ring that facilitates observation of the samples and definition of the OCT scan direction/area.

The operating principle of OCT is based on white light interferometry similar to a Michelson type interferometer (Fercher et al., 2003) (Fig. 1), where broadband low coherent light from the SLD is split by a beam splitter into two partially coherent beams, a reference beam and a sample beam. The sample beam penetrates the object along the *z*-axis, while the reference light is reflected by a retro-reflecting mirror within the imaging probe generating a reference beam of known path length. In a 1-dimensional A-scan, the sample beam interacts with the coral tissue and light is reflected from different microstructural features along the *z*-axis. The locally reflected light is collected and combined with the reference beam, generating a characteristic interference pattern. In time-domain OCT, the reference mirror is rapidly scanned along the *z*-axis in order to vary the optical path length and thus imaging a particular depth of interest. In spectral-domain (SD) OCT, which was applied here, no mechanical scanning of the reference arm is needed and instead the cross-spectral density is measured by a spectrometer that characterizes the unique phase delay of each wavelength, which significantly improves the imaging speed of the OCT system. An A-scan’s interferometric signal from each *x*,*y* position is converted into a signal of reflectance as a function of depth (z) with 4.5-μm axial resolution, which allows separation of the different coral tissue layers and skeleton scatter (Wangpraseurt, 2016). To generate 2-D transects (B-Scans) and 3-D stacks of transect (C-scans), a galvanometer system inside the imaging probe moves the A-scan sample beam along the *x* and *y* axis. The build-in USB camera in the OCT scan head enabled us to define exact scan positions, transects and areas relative to the visible coral structure.

The SD OCT system used here allows for video rate scanning, with a maximal A-scanning rate of 36 kHz given there is sufficient contrast in the sampled OCT-signal. Conventionally, the SD OCT data is described as a signal to noise ratio (in decibel, dB).The generated SD OCT data represents a relative measure of the locally backscattered light and carries quantitative optical information that can be used to calculate the scattering coefficient of a biological tissue (Faber et al., 2004). However, the characterization of tissue optical properties necessitates a careful calibration of the OCT detector optics and measuring setup. Calibration of the OCT data in absolute units (i.e. reflected power) is rarely performed in OCT studies, where the primary aim is the visualization of structural features. In this study, the OCT data is thus shown in dB and visualization was aided through false color coding and gray scaling of the OCT dB signal (see below for details).

### 2. 2. Experimental procedures

#### 2.2. 1 Sampling of corals

Sun-adapted corals were collected from shallow waters (<2 m depth) on the reef flat of Heron Island, Great Barrier Reef, Australia (152°06’E, 20°29’S). Fragments of the corals *Acropora millepora, Acropora aspera, Acropora pulchra, Cyphastrea serailia, Favites abdita, Favites flexuosa, Goniastrea aspera, Lobophyllia corμmbosa, Goniastrea australiensis, Pocillopora damicornis* (brown and pink morph), and *Stylophora pistillata* were selected; coral identification was done according to Veron (2000). The coral species were chosen to cover a wide range of different surface structures and skeletal architectures as well as different types of tissue thickness and pigmentation e.g. due to the presence of GFP-like coral host pigments. After collection, colonies were fragmented into smaller pieces of a few cm^−2^ in size, mounted onto microscope slides and photodocumented. All corals were kept under natural light conditions in outdoor aquaria at the Heron Island Research station supplied with a constant flow of fresh seawater.

#### 2.2.2 OCT imaging setup

Coral OCT measurements were done indoors under defined light conditions provided by a fiber-optic halogen lamp (KL-2500LCD, Schott GmbH, Germany) with individual coral samples positioned in a custom-made black acrylic flow chamber connected to a 10 L water reservoir with fresh aerated seawater (Wangpraseurt et al., 2012a). The flow velocity was about 0.5 cm s^−1^ with a constant water temperature of ~24°C and a salinity of 33-35. To optimize the signal to noise ratio in the measurements, the corals were positioned within the flow chamber such that only a small amount of water (<3 mm) was on top of the coral surface, which minimized water absorption of the 930 nm sample light beam; thinner water layers resulted in optical artefacts due to air-water reflections. It was also important to minimize flow chamber vibrations, which otherwise led to interference artefacts in the OCT scans.

Measurements on intact corals were performed by selecting a region of interest (ROI) using the image generated by the USB camera of the OCT system. The USB camera image was first brought into focus using the manual focusing stage. The reference length of the scan head was then adjusted until the OCT signal was maximized within the uppermost 30% of the image field. OCT scanning was performed in B-Scan (cross sectional) and C-scan (3-D rendering) mode using an A-scan averaging of at least 10 scans with an A-scan rate of 36 kHz. Fast measurements of single B-scans were also performed to follow dynamic changes in coral tissue structure due to light-induced tissue contraction (Wangpraseurt et al., 2014). For this, a *Favites abdita* fragment was kept in darkness in order to lead to a complete relaxation of the oral polyp tissue, which could be followed by OCT scanning in darkness. The polyp was then illuminated by a fiber optic tungsten halogen lamp (see above) providing a downwelling photon irradiance (PAR, 400-700 nm) of *Ed* = 3000 μmol photons m^−2^ s^−1^, while the tissue contraction was followed in OCT B-scan timeframe mode, recording subsequent B-scans at a frame rate of 0.7 seconds. Additionally, we tested the suitability of OCT to image the exposed skeleton by removing the coral tissue with an air-gun and repeating OCT measurements under the same underwater conditions. For coral skeleton imaging, it was important to release any air trapped between skeleton ridges upon immersion by carefully tapping the skeleton and brushing its surface underwater with a paintbrush.

### 2.3 Image analysis

Visualization of OCT B‐ and C-scans was done with the manufacturers imaging software (Thorimage 4.2; Thorlabs GmbH, Luebeck, Germany) using the in-built thresholding functions. OCT C-scans were saved as ‘raw + processed’ data in the software and were then converted to 32-bit grayscale multiple tiff stacks using a custom-made macro written in ImageJ (Wagner et al., 2010). Post-processing was performed in a Fiji installation of ImageJ (Schindelin et al., 2012) as described below.

#### 2.3.1 OCT based imaging of GFP-like coral host pigments

A protocol was developed for the non-invasive characterization of GFP chromatophore abundance and size by OCT, as preliminary observations revealed that OCT is capable of imaging and identifying the highly reflective chromatophores of GFP-like host pigments in the coral *Favites abdita* (Lyndby et al., 2016). For this, OCT C-scans were rendered in Thorimage 4.2, and a manual threshold analysis was performed in monochrome modus to maximize the image contrast (in terms of the OCT dB signal) between GFP chromatophores and the surrounding tissue. The rendered OCT 3-D scan was exported as a JPG image with a *z*-axis orientation of 0° (i.e. top view) and imported into ImageJ. For the investigated coral polyps, OCT B-scans revealed that the chromatophores were exclusively located as a single layer of granules within the upper oral tissue layer allowing for a particle density estimate based on a single rendered JPG file.

Three 1 mm^2^ regions of interest (ROI) were manually selected within image areas that showed maximal signal/noise ratios. For each ROI image, brightness and contrast were adjusted, followed by 8-bit conversion and binarisation via the Otsu segmentation algorithm (Zhang and Hu, 2008) plugin in ImageJ. The Otsu automatic thresholding algorithm minimizes the intra-class variance between black and white pixels (Zhang and Hu, 2008) while ensuring no significant loss of granule features. The binarised image was then analysed for granule density using the ImageJ automated particle analyser plugin. Granule detection was optimized through size boundaries (pixels^2 = 7.5 − 8) and hole filling, while the image edges were excluded. The particle analyser calculated the chromatophore number per projected surface area, chromatophore size (in μm projected surface area), as well as the % surface cover by chromatophores. Additionally, the average maximal diameter of GFP chromatophores was estimated from OCT B-scans, i.e., tissue cross-sections, using the manual thresholding function of Thorimage 4.2.

#### 2.3.2 Linear velocity of tissue contraction

The linear velocity of coral tissue contraction was calculated from recorded time-series of B-scans using an automated single-particle tracking algorithm, TrackmateJ (Tinevez et al., 2016). The frame sequence was first exported from Thorimage 4.2 in RGB mode and converted to 8-bit grayscale images in ImageJ. For each frame, a white circle (width = 25 pixels) was added adjacent to the tissue surface to aid in boundary detection (see Movie 3). The tissue surface contraction was tracked in the constant speed tracking mode of TrackmateJ using an estimated blob diameter of 100 pixels and a threshold of 2 (Tinevez et al., 2016). The maximal linear velocity was calculated in μm s^−1^ as the running average of the vertical displacement over 3 subsequent frames (about 2.2 seconds) for each frame.

#### 2.3.3 Coral tissue surface rendering and surface area estimates

The coral tissue surface rendering and surface area analysis involved image thresholding and binarisation, development of a topographic height map followed by a gradient analysis. Extracted OCT C-scans (32-bit grayscale tiff stacks) were first adjusted for image brightness and contrast following conversion to 8 bit in ImageJ. The stacks were then binarised using the Otsu automatic thresholding algorithm (see above). Subsequently, remaining noise, i.e., white pixels not related to the coral surface, was removed using the remove outlier function of ImageJ. The images were cropped such that maximal tissue surface height equaled *z*=0. A depth coding algorithm (Wagner et al., 2010) converted the binarised tiff stacks to a continuous height function *z*(*x,y*), where each *z* value was assigned a pseudo color-coded pixel value. The generated height map was visualized as a 2-D surface plot in order to reconstruct the surface topography of the OCT C-scan.

The surface area of the reconstructed surface topography was then calculated using SurfCharJ, an ImageJ plugin originally developed for paper structure analysis (Chinga et al., 2007). SurfCharJ estimates surface roughness and surface area by performing a gradient analysis that measures the intensity and orientation of local surface structures of the topographic image. The orientation of a surface in 3-D space can be described by the direction of the outward surface normal vector, 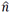 (Chinga et al., 2003). In polar coordinates 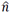 is given in terms of the polar angle, *θ*, and the azimuthal angle, *ø*, by

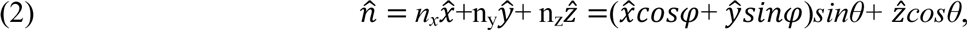

where 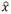, *ŷ* and 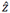 are unit vectors of the respective axes. The topographic height data relate to the coordinate angles (*θ, ϕ*) by:

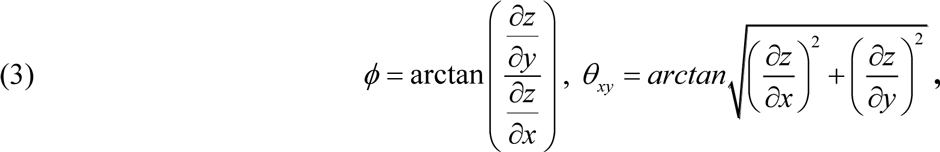

where *z* is the surface height. Equation 5 and 6 show how the angles are related to the gradients of the height data in the topographic map. The gradients are then estimated using a series of imaging processing algorithms that are described in further detail in Chinga et al. (2007). Briefly, the gradient analysis calculates the orientation of the local surface relative to the coral surface, i.e., the polar image, and the direction of the local surfaces relative to the coral tissue plane, i. e., the azimuthal image. The coral tissue surface area (SA) is then calculated based on the gradient analysis as:

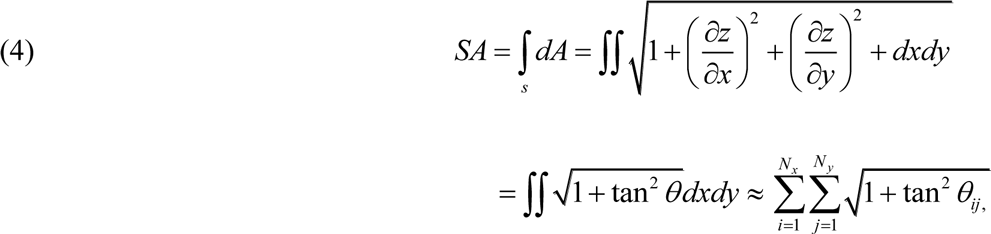

The coral tissue surface area could thus be calculated as the ratio between the actual coral tissue surface area (in μm^2^) relative to the planar area imaged with OCT (in μm^2^).

### 3. Results and Discussion

#### Basic microstructural features of coral tissues

OCT is a non-invasive high resolution imaging technique that has been widely used in the biomedical field, especially in dermatology and ophthalmology (Huang et al., 1991). Recently, OCT imaging has also been applied in the environmental sciences in order to understand the structure and function of biofilms (Wagner et al., 2010), higher plants (Hettinger et al., 2000; Kutis et al., 2005), aquatic vertebrates and invertebrates (Bellini et al., 2014; Speiser et al., 2016). We present the first application of OCT for studying the *in vivo* tissue and skeleton structure of tropical corals.

The tissues of all 11 species of investigated corals, spanning a wide range of morphologies and tissue thickness, could be imaged with excellent optical contrast (Fig. 2). Signal contrast in OCT depends on the optical scattering properties of the investigated material, where light scattering due to refractive index mismatch creates optical signal and thus good image contrast, while highly absorbing media result in poor images (Fercher et al., 2003). Recently, it has been shown that coral tissues exhibit strong light scattering (Wangpraseurt et al., 2014; Lyndby et al., 2016) with scattering coefficients similar to human and plant tissues (Wangpraseurt et al., 2016). Therefore OCT was ideally suited to resolve microstructural features of coral tissues (Fig. 2, 3).

**Fig. 2.**
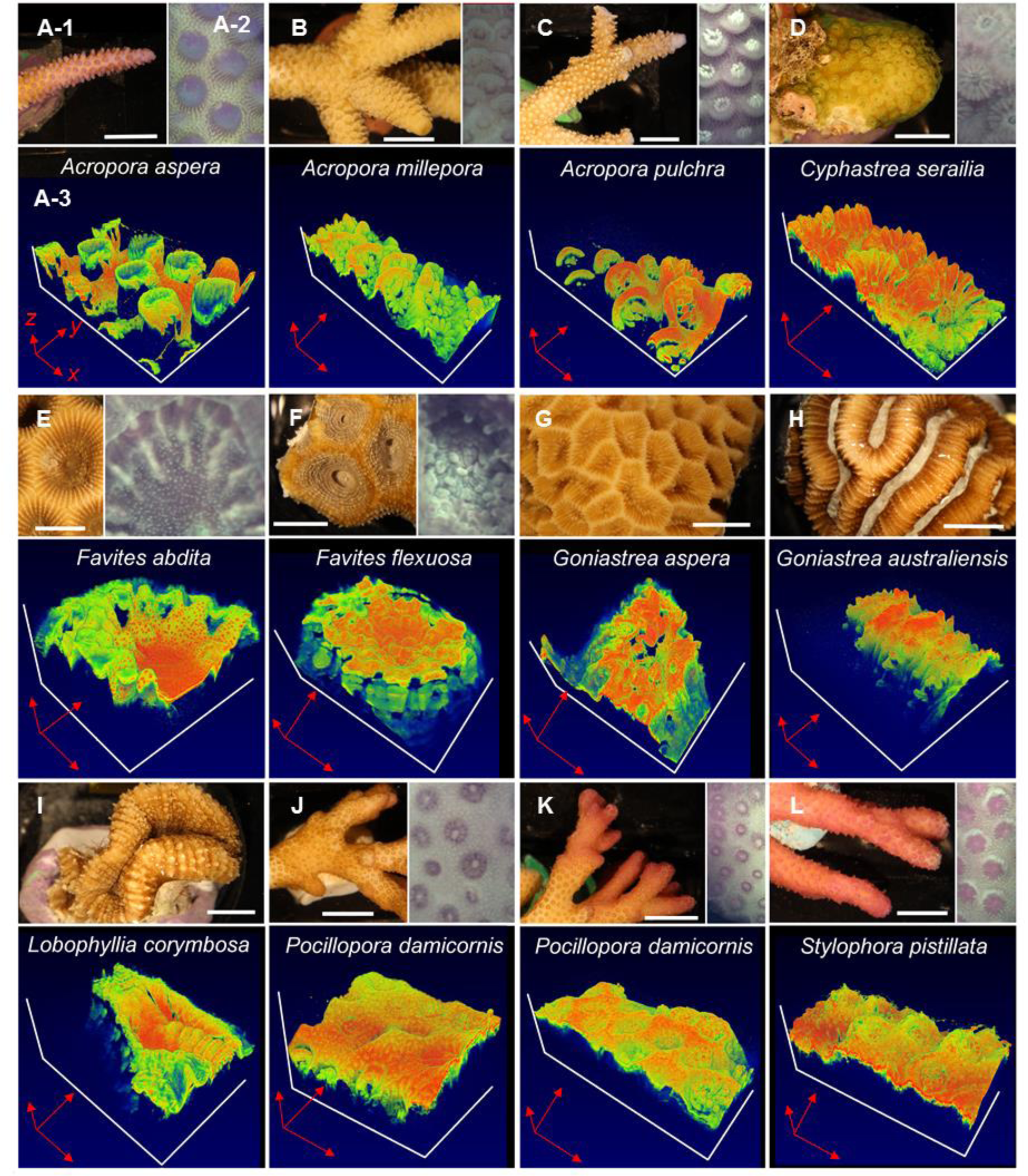
A-M. Three-dimensional OCT imaging of coral microtopographic diversity. A-1. Macrostructure of an *Acropora aspera* branch (white scale bar = 1 cm). **A-2** Close-up, top view of *Acropora aspera* tissue microstructure photographed with the USB camera of the OCT system. The photographed tissue surface area corresponds to the area imaged with OCT shown in panel **A-3.** The three-dimensional OCT scan is shown in *x,y,z* dimensions. Red arrows are 2 mm in length for *x,y* and 1 mm in length for the *z* dimension. The false color coding represents the intensity of the uncalibrated OCT signal, which was manually adjusted for to optimize visualization of structural features for each scan (see methods for details). Note that panel number labeling is only shown for panel **A** (for clarity) and that no USB camera image was taken for some coral species as OCT scanning was performed in darkness to ensure tissue relaxation.

**Fig. 3:**
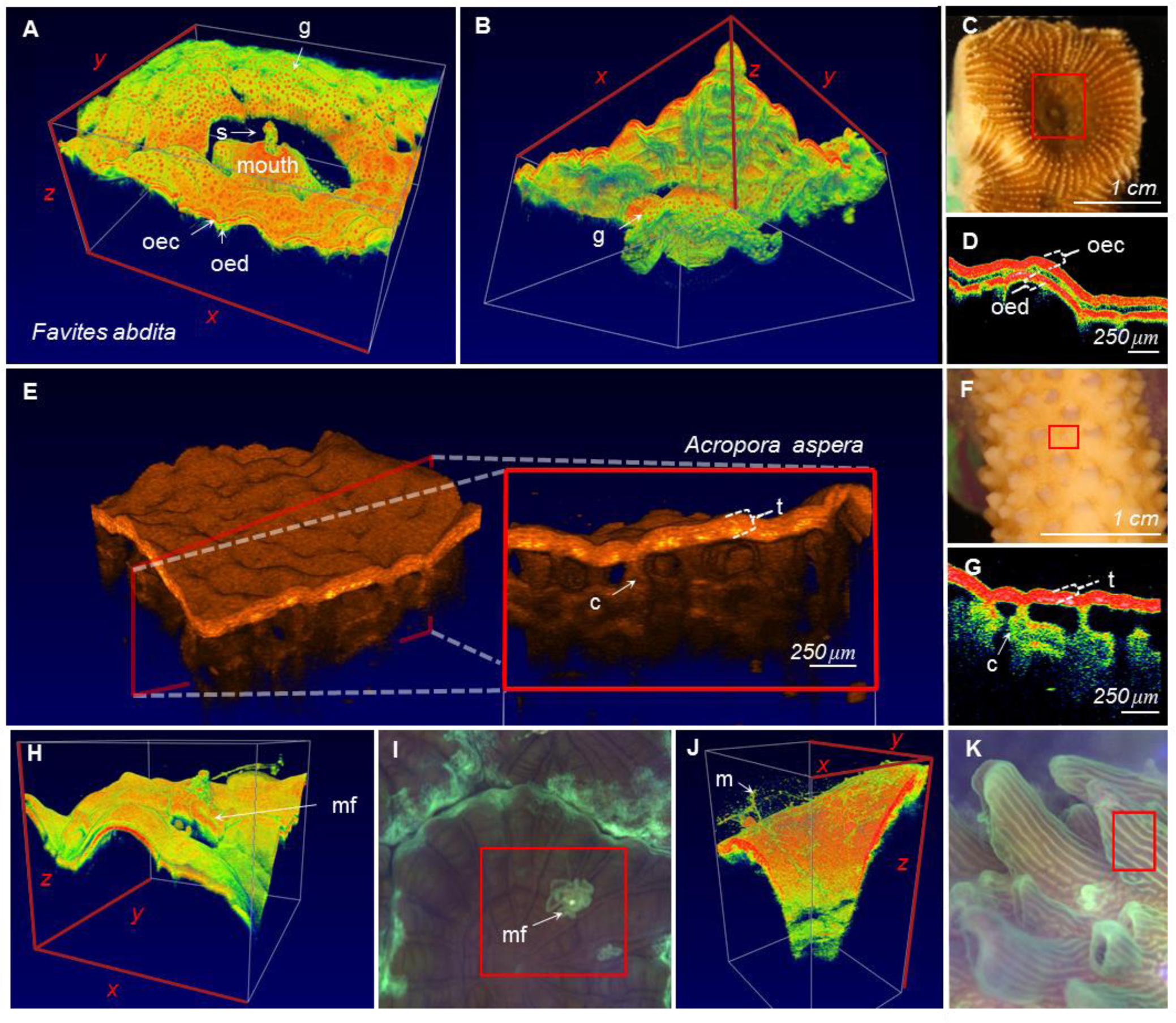
Microstructural imaging of coral tissues using OCT. A-D) *Favites abdita*. **A)** Three-dimensional rendering of a single polyp. The rendering visualizes the convoluted surface topography and polyp mouth and identifies the oral ectoderm (oec), the oral endoderm (oed), GFP granule containing chromatophores (g) as well as sediment ingestion (s). **B)** rear view of panel **A**. The field of view of the OCT scan was *x*=5.25 mm, *y*=4.4 mm and *z*=2.8 mm. Video animation of the 3D rendering can be found in the supplementary information (**Movie 1**). **C)** image of coral polyp. The red square illustrates the approximate area imaged with OCT. **D)** close up of tissue arrangement in cross-sectional OCT B-scan mode**. E-G)** *Acropora aspera*. **E)** three dimensional inlet shows representative *en face* plane of a coenosarc area. The scan identifies the entire tissue (t) and skeletal channels (c). **F)** photograph of live coral, red square shows approximate OCT scan area, **G)** close up showing the channel like structures. **H)** three dimensional rendering of mesenterial filaments (mf) based on imaged area (red square) in panel **I**. (*x=*3.0 *y*=3.4, *z=*2.8 mm) **J)** three dimensional rendering of coral mucus (m) based on imaged area (red square) in panel **K** (*x=*1.1 *y*=1.45, *z=*2.8 mm). Video animation of coral mucus movement can be found in the supplementary information **(Movie 2**) False color coding blue to red (highest signal in red) and orange tones (highest signal in brightest tones) was manually adjusted to optimize image contrast.

Close-up OCT scans of the oral disc of the massive thick-tissued coral *Favites abdita* revealed a highly heterogeneous coral tissue surface with tissue lobes surrounding the immediate mouth opening (Fig. 3 a-d). OCT B-scans revealed the oral ectoderm which was ~ 90 μm thick followed by a thin (~50 μm layer) of low OCT signal which likely relates to the mesoglea (Fig. 3D, Fig. S1). The oral endoderm was also visible and at least 30 μm thick (Fig. S1). Along the A-scan axis beyond the oral endoderm, the OCT signal was quickly attenuated and imaging penetration depth was around 500 μm from the coral surface. Thus for the thick-tissued *F. abdita*, the OCT imaging was limited to the upper tissue layers (Fig. 3b, d).

In contrast, imaging of the thin-tissued *Acropora aspera* generated higher axial penetration depths of up to 1 mm, visualizing the entire coral tissue and the underlying skeleton (Fig. 3 e-g). The coral tissue of *A. aspera* appeared as one continuous tissue layer due to the steady linear attenuation of the OCT signal along the A-scan axis within the coral tissue (Fig. 3D and Fig. S1). OCT signal attenuation within *A.aspera* was lower compared to *F. abdita*, which allowed for imaging the aragonite skeleton below the animal tissue layer (Fig. 3E, G). The OCT scans further revealed that the tissue was connected to the skeleton via channel-like ‘pillars’ that were on average 20-50 μm in diameter (see ‘c’ in Fig. 3G). Scanning electron microscopy of dead coral skeletons showed the presence of such skeletal extrusions of 40-100 μm width in *Acropora aspera* (Brown et al., 1985). OCT now allows for imaging the tissue-skeleton interface of live corals without the need for tissue removal and/or decalcification, thus facilitating the *in vivo* investigation of skeletal structures and growth e.g. through repeated monitoring of defined tissue and skeleton section of the same individual (Tambutté et al., 2007). Additionally, OCT can also be used to only image the exposed skeleton with good optical contrast (Fig. S3) and might thus provide a suitable alternative to e.g. 3-D laser scanning. A thorough analysis of coral skeleton morphology with OCT was, however, beyond the scope of the present study.

The vertical penetration depth and image contrast for the employed OCT system is affected by NIR (930 nm) attenuation, primarily due to absorption in water (Kirk, 1994) and multiple scattering in coral tissue and skeleton (Wangpraseurt, 2016; Wangpraseurt et al., 2012a). The OCT signal originates from directly backscattered photons, while multiple scattering reduces image resolution and contrast with increasing sampling depth, where the operational limit for the OCT imaging penetration depth is affected by the scattering mean free path, i. e., the inverse of the reduced scattering coefficient *μ*_*s*_’. The reduced scattering coefficient is:

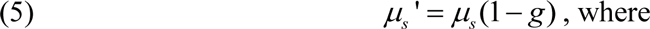

*μs* is the scattering coefficient and *g* is the anisotropy of scattering, i.e. the directionality of the scattering (Fercher et al. 2003). Most biological tissues are highly forward scattering (*g* ~ 0.9; Jacques, 2013) and the operational penetration depth of OCT imaging typically is ~3-5 times the scattering mean free path (Fercher et al. 2003). Assuming *g*~0.9 and a coral tissue scattering coefficient *μs* of ~100 cm^−1^ in dense Faviid polyp tissue (Wangpraseurt *et al.* 2016), we would thus expect a depth limit for OCT imaging of ~300-500 μm, which fits our experimental observations in *Favites abdita* (Fig. 2, Fig. S1). The operational OCT imaging depth was larger in Acroporids and other branched corals, which might reflect a lower scattering coefficient and/or lower overall attenuation of the 930 nm OCT probing light in such thin-tissued corals. A more precise quantification of how different corals and different coral structures affect the operational depth of OCT awaits further quantification of the inherent optical properties of corals (Wangpraseurt et al., 2016). Provided that the depth dependent light collection efficiency of the OCT system is known it is possible to extract quantitative estimates of the scattering coefficient (*μs*) and anisotropy of scattering (*g*) based on the local reflectivity (*φ*) and the OCT signal attenuation (*μ*) (Faber et al., 2004; Levitz et al., 2010), but such approaches were beyond the scope of the present study.

#### Imaging coral tissue and mucus dynamics

Research in coral reef science has been dominated by studies on the response of corals to environmental stress including global warming, ocean acidification, pollution and diseases (Hoegh-Guldberg et al., 2007; Hughes et al., 2003). Now OCT imaging can potentially facilitate the early detection of ecophysiological stress through microscale visualizing of behavioral modifications *in vivo.* For instance, mesenterial filaments can be expelled from the polyp stomach and extruded to the coral surface e.g. in response to temperature stress (Brown et al., 1995) and space competition with neighboring corals and predators (Nugues et al., 2004). A recently, developed underwater microscope was able to record mesenterial filament extrusion at high spatial and temporal resolution (Mullen et al., 2016). Likewise, OCT was capable of following and visualizing the extrusion of such filaments through temporary openings with microscale spatial resolution (Fig. 3H-I). The good visibility of mesenterial filaments is likely related to their high lipid content (Brown et al., 1995) creating a sufficiently strong refractive index mismatch with the surrounding medium to result in excellent contrast and visualization through OCT (Fig. 3H).

Coral mucus is fundamental to the energetics of coral reefs and mucus can comprise up to 20-45% of the energetic demands of corals (Wild et al., 2004). Mucus excretion is affected by e.g. temperature and light stress, bacterial infection, particle sedimentation and coral exposure to air (Brown and Bythell, 2005; Philipp and Fabricius, 2003). There has been substantial interest in imaging and quantifying coral mucus, but visualization has been hampered as it is almost invisible in conventional light microscopy, and mucus quantification has thus mainly relied on volumetric bulk measurements (Koren and Rosenberg, 2006; Wild et al., 2005). Mucus is essentially a hydrated gel (Brown and Bythell, 2005) and quantification via histology suffers from artefacts during sample preparation due to dehydration and subsequent shrinkage (Jatkar et al., 2010). Digital holographic microscopy (DHM) was recently used for visualizing the *in vivo* mucus secretion in cold-water corals (Zetsche et al., 2016) and this novel technique seems well suited for similar studies on symbiont-bearing corals. Here we show that OCT is a suitable alternative to DHM as it was able to resolve the string and sheath-like mucus structures adhering to the coral surface (Fig. 3J-K, Fig. S2). By exposing the coral *Acropora aspera* to air for about 30 seconds, we were able to stimulate excessive mucus production. Due to the rapid imaging acquisition of OCT, mucus excretion and transport can now be quantified and followed in real time (Fig. S2 and Movie 2). However, it is important to note that any light scattering particles trapped within the mucus will also be imaged by OCT, and thus care must be taken when interpreting such image-based data on mucus secretion in quantitative terms.

OCT is not a suitable tool to image microalgal loss during coral bleaching (Fig. 4). The optical properties of *Symbiodinium* are dominated by light absorption (Teran et al., 2010), although microalgal cells do also scatter light (Morel and Bricaud, 1986). We imaged a bleached polyp of the coral *Pocillopora damicornis*, revealing no major differences in the OCT signal generated from the polyp as compared to healthy tissues (Fig. 4a-d). Sectioning of the coral polyp revealed clearly the stomach tissues of the polyp (Fig. 4d). The good visibility of coral tissues independent of *Symbiodinium* density, suggests that OCT can be used to detect early sub lethal signs of environmental stress such as changes in tissue thickness prior to coral bleaching (Ainsworth et al., 2008). Hitherto, changes in coral microstructure upon temperature stress have primarily been assessed with invasive imaging techniques that require tissue sectioning such as conventional light microscopy or scanning electron microscopy (Ainsworth et al., 2008; Glynn et al., 1985). In contrast, the non-invasive OCT approach now allows rapid and repeated *in vivo* monitoring for early signs of stress responses in intact corals.

**Fig. 4:**
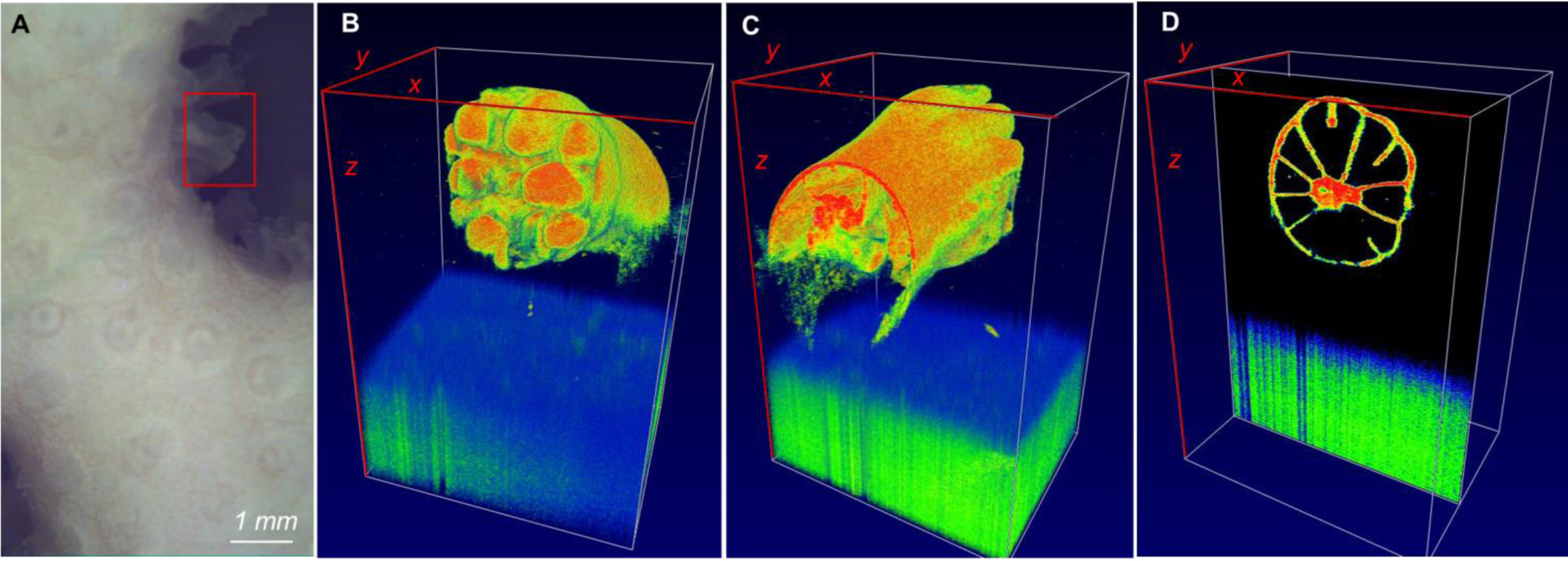
OCT imaging of a single bleached coral polyp. **A)** The image shows the bleached surface of the coral *Pocillopora damicornis*. The red square corresponds to the area scanned with OCT. **B-C)** Three dimensional rendered coral in volume view, and **D)** in sectional view. The field of view of the OCT scan was *x*=1.2 mm, *y*=9.45 mm and *z*=2.8 mm.

#### Visualization of coral host pigments

The coral animal host has several potential means to protect its photosymbionts from excess radiative stress that can otherwise cause photoinhibtion (Baird et al., 2009). The coral animal synthesizes various fluorescent (Alieva et al., 2008) and non-fluorescent pigments (Dove et al., 2008) as well as myscroporine-like amino acids (MAA) (Baird et al., 2009; Shick and Dunlap, 2002) all of which are thought to have photoprotective functions (Dunlap and Shick, 1998; Lyndby et al., 2016; Salih et al., 2000). GFP-like host pigments are homologous to the well-known green fluorescent proteins that are now widely used as a fluorescent markers in life sciences. There thus exists a major body of research dealing with the structure and function of GFP-like pigments in corals (e.g., Alieva et al., 2008; Oswald et al., 2007; Salih, 2012). GFP-like pigments can be loosely dispersed in the coral animal tissue or occur in pigment granules (about 1μm in size) that can aggregate to a chromatophore system (Salih et al., 2003; Salih et al., 2000; Schlichter et al., 1994). *In vivo* visualization of GFP-like host pigment granules has been achieved by confocal microscopy using their natural fluorescence to create image contrast (Salih et al., 2000). However, in addition to their fluorescent properties, GFP-like pigments are also highly scattering (Lyndby et al., 2016). In this study, we demonstrated that OCT was able to distinguish and identify chromatophores containing GFP-like host pigment granules with good contrast from the remaining coral tissue, due to their strong light scattering properties causing strong OCT signals (Fig. 5a-c).

**Fig. 5.**
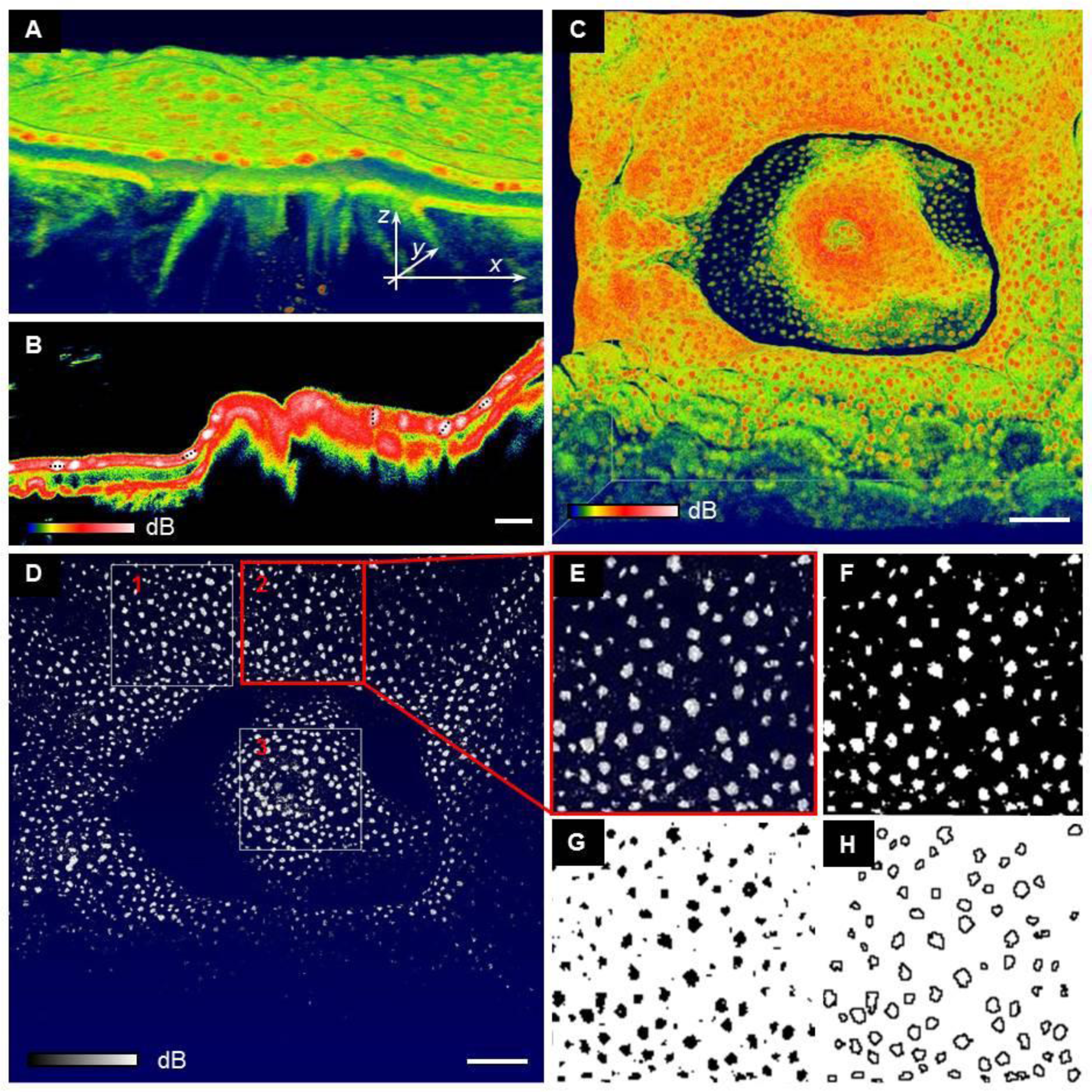
Imaging and segmentation of GFP containing chromatophore system. **A)** close-up of a 3D rendering of the *Favites abita* polyp shown in Fig. 1E. Highest OCT signal intensity (in red) reveals chromatophores. Length of arrows equals 500 μm. **B)** cross-sectional OCT B-scan through the polyp mouth. Chromatophore visualisation (in white) was enhanced through narrowing the threshold of the OCT signal. Dotted black lines show examples of maximum crhomatophore diameter. Scale bar=100 μm **C)** Top view of 3D rendering of coral polyp in false color mode and **D)** in black and white color mode. Scale bar = 500 μm. Regions of interest (labelled 1-3) of 1 mm^2^ quadrats were selected for chromatophore analysis. **E-H)** Example of segmentation steps to yield physical characteristics of chromatophores**. E)** Close-up of region of interest 2, **F)** brightness and contrast adjusted image, **G)** binarised image using the Otsu algorithm and **D)** detected particles (see methods for details).

For the investigated *Favites abdita* coral, GFP chromatophores were exclusively located in the upper ectoderm (Fig. 5a,b), where the density of GFP chromatophores within the coral polyp tissues was on average 97 (±15) granules per mm^2^ of projected two dimensional surface area, which translated to a surface cover of 13% (±4%) within the imaged polyp tissue. The maximal diameter of the chromatophores was on average 85 μm (±13.8) as assessed through OCT B-scanning. Size-frequency distributions calculated a median chromatophore size of 1.0-1.5 x 103 μm^2^ per two-dimensional projected surface area (Fig. 6). Previous studies have characterized GFP granules and reported on the existence of the chromatophore system (see e.g., Salih et al., 2000; Schlichter et al., 1985), but a quantification of the abundance and size structure of chromatophores over larger coral tissue areas has hitherto been lacking. Photomicrographs of thin histological sections shown in Schlichter et al. (1994) revealed the presence of chromatophores roughly about 60 μm in diameter. Likewise, confocal microscopy stacks showed the presence of chromatophores within the same size category (~50-100 μm) (Salih et al., 2000; Salih et al., 2004). Thus although there is no comparable size distribution estimate on the reported chromatophore system, our OCT-based size estimates fall within the range reported in the earlier studies. While OCT is a good method to rapidly assess the basic properties and *in vivo* abundance of chromatophores within larger tissue areas without the need for physical preparation, more diffusely incorporated GFP pigments were not clearly discernible and the two morphotypes of *Pocillopora damicornis* (brown, non-fluorescent vs pink, fluorescent) did not reveal characteristic differences in their OCT signal (Fig. 1J and K).

**Fig. 6.**
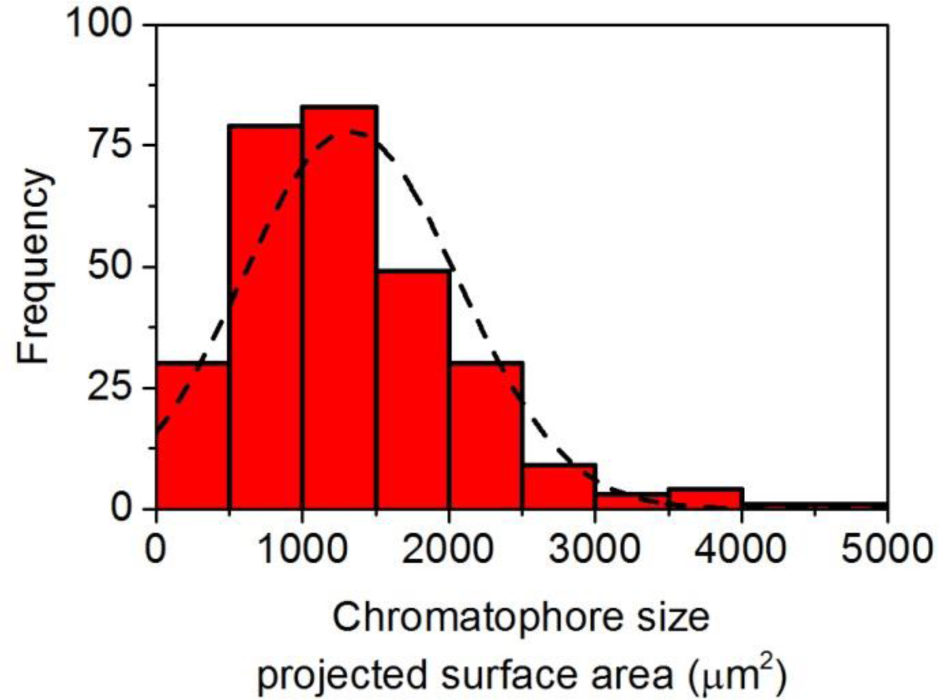
Size-frequency distribution of GFP containing chromatophore system of *Favites abdita*. Chromatophore size is calculated in relation to the projected two dimensional surface area (μm^2^). Chromatophore identification is based on image analysis shown in Fig. 5.

#### Coral tissue plasticity

Although corals appear to be static on the macroscale, coral tissues are flexible on the micro to cm scale, where coral tissues can re-arrange due to e.g. physical force, behavioral control or illumination (e.g. Mullen et al., 2016). Massive thick-tissued corals respond to excess illumination via tissue contraction (Levy et al., 2003), which can increase tissue surface reflectivity and affect the internal tissue light microenvironment (Wangpraseurt et al., 2014). However, a precise quantification of such tissue plasticity has been lacking. We therefore traced the linear velocity of tissue contraction using rapid OCT B-scanning (Fig. 7). Upon illumination, a *Favites abdita* polyp accelerated within 3-5 seconds to a maximum linear tissue movement velocity of ~120 μm s^−1^, whereafter the velocity decreased linearly until movement stopped within ~20 seconds. In the current example, we captured B-scans of 4 mm wide x 2.8 mm deep vertical sections of intact coral tissue and skeleton at a framerate of 0.7 seconds (Movie 3). Reducing the spatial resolution and field of view to dedicated areas would enable coral movements to be followed in real time, i. e., at video rate resolution (Blauert et al., 2015)

**Fig. 7.**
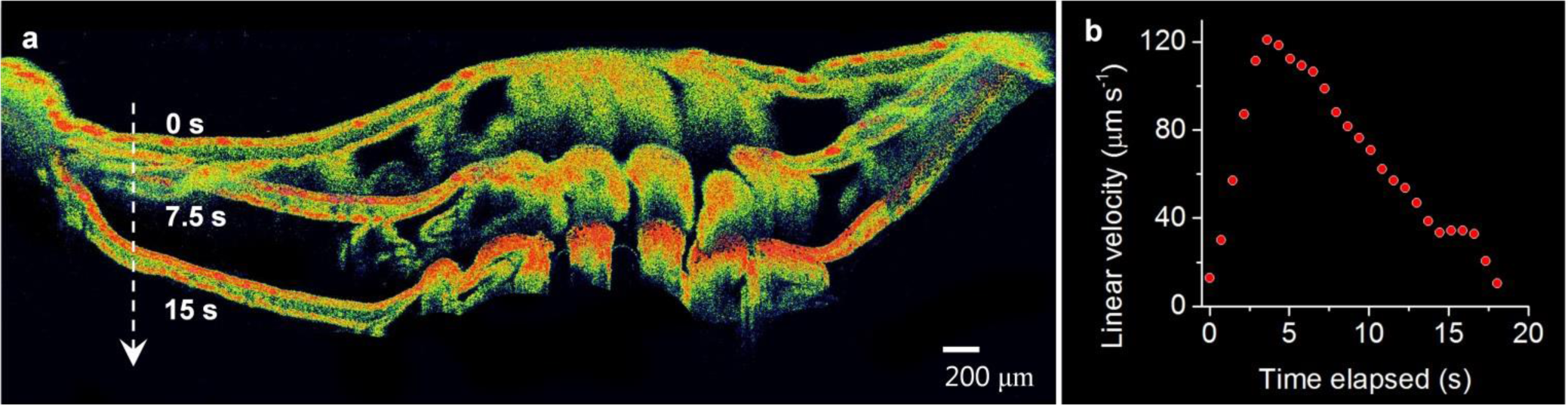
*In vivo* cross-sectional tracking of tissue movement. **a)** Sequence of OCT B-scans (false color mode) performed after illuminating the coral *Favites abdita*. The three scans show the coral polyp in an expanded state in darkness (0 seconds) and after 7.5 seconds and 15 seconds of high light illumination. The 3 images are super-imposed for visualizing tissue plasticity (see supplementary information **Movie 3** for real time video of tissue contraction). **b)** Estimated maximal linear velocity of tissue movement calculated as the running average of the vertical displacement of a white marker (see methods) towards the center of the image over 3 frames (about 2.2 seconds).

Gas exchange is regulated across the tissue-water interface and thus partially controlled by the area of exposed tissue surface controlled by tissue contraction and expansion (Patterson, 1992; Patterson and Sebens, 1989). Corals thus have the capacity to alter gas exchange dynamics through significantly changing their exposed surface area. *Symbiodinium* light exposure will also be affected by changes in tissue contraction and expansion (Wangpraseurt et al., 2014). Changes in tissue surface structure are thus likely to affect tissue surface and subsurface scattering as well as the optical path length between coral tissue surface and *Symbiodinium*. OCT provides a method to quantify such dynamic changes in surface structure and tissue density. Surface area is thus an important metric in coral biology research, where metabolic rates and e.g. *Symbiodinium* cell numbers are often normalized to coral surface area. Coral surface area has mainly been estimated from coral skeleton architecture through e.g. wax dipping and foil wrapping techniques (Marsh, 1970; Stimson and Kinzie, 1991) or x-ray scanning based computer tomography (μCT; Naumann et al., 2009). Direct surface area quantification on intact corals used has also been attempted with photogrammetry (Courtney et al., 2007; Holmes, 2008; Naumann et al., 2009), but these techniques have been established for surface area characterization on centimeter and meter scales, largely neglecting the micro-architectural complexity and plasticity of coral tissue surfaces (Fig. 2). Here we developed an OCT-based method to rapidly quantify coral surface area for a given tissue state (see Methods). We calculated the change in three dimensional tissue surface area of the coral *Pocillopora damicornis* upon tissue contraction (Fig. 8) as the ratio of the three dimensional surface area normalized to the two dimensional projected surface area. Our calculations showed an approximate doubling (2.14 times) of relative tissue surface area upon tissue expansion from 2.2 (contracted) to 4.5 times (expanded) in the coral *Pocillopora damicornis*. This illustrates the importance of taking such dynamic changes in tissue structure into account e.g. when normalizing rates of photosynthesis or respiration to coral surface area. A detailed comparison of OCT-based surface area determination with existing methods was, however, outside the scope of the present study and is now underway.

**Fig. 8.**
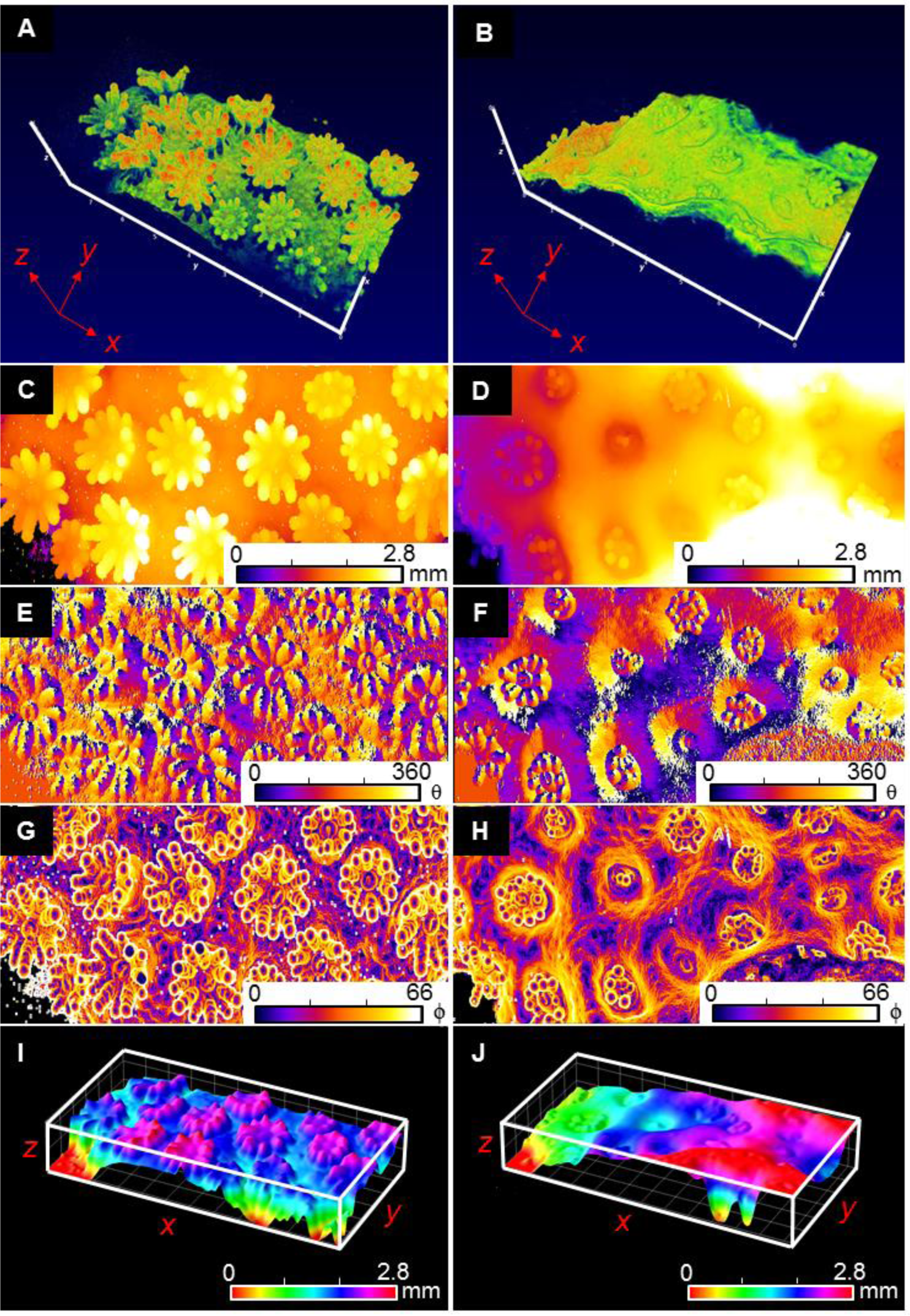
OCT-based surface area characterization of expanded (left panel) vs contracted coral tissue (right panel). **A, B)** Three dimensional OCT scan. (*x,y,z* scale bar = 1mm). **C, D)** Topographic height map (z=0-2.8 mm in false color coding). **E, F)** The gradient analysis (see methods) calculated the azimuthal angle θ showing the surface direction from 0 to 360°, and **G, H)** the polar angle ø showing the surface orientation from 0 to 66°. **I, J)** The reconstructed three dimensional tissue surface (see methods). The surface height *z* is shown in false color coding for *z*= 0-2.8 mm. The field of view of the OCT scans was *x*=3.4 mm, *y*=7.7 mm and *z*=2.8 mm.

### Conclusion

Optical coherence tomography is a non-invasive imaging technique that can easily be applied for studying the structure of tissues and skeleton of intact living corals *in vivo*, where the use of NIR avoids any actinic effects on the coral host or photosymbionts. Coral tissue and skeleton microstructure can be imaged with excellent optical contrast and OCT further allowed for the characterization of coral tissue surface area, distribution and abundance of GFP-like pigment granules, and quantification of coral tissue contraction at a hitherto unreached spatio-temporal resolution on living corals. OCT is also well suited for monitoring early responses of corals to environmental stress in the form of mucus excretion and mesenterial filament extrusion. This first application of OCT in coral reef science also points to several future developments. For instance, biomedical OCT analysis has developed image analysis tools that allow for advanced image correction for optical artefacts such as speckle noise and refraction, as well as for dedicated segmentation of microstructural features (Kafieh et al., 2013), and implementation and optimization of such tools for coral OCT analysis seems promising. The same holds true for the use of calibrated OCT measurements for extracting inherent optical properties (Faber et al., 2004; Levitz et al., 2004) such as the scattering coefficient of coral tissue and skeleton. OCT systems are compact, robust and can easily be incorporated into typical experimental setups for ecophysiological studies of corals, and inclusion of OCT in such studies can provide key information about coral structure such as e.g. *in vivo* quantification of tissue surface area dynamics under different experimental treatments.

## Conflict of Interest

*The authors declare that the research was conducted in the absence of any commercial or financial relationships that could be construed as a potential conflict of interest*.

## Author Contributions

DW and MK designed study. DW and CW performed measurements. SLJ and MW provided analytical tools. DW analyzed data with input from SLJ, MW and MK. DW and MK wrote the article with editorial input from all co-authors.

## Funding

This study was funded by a *Carlsberg Foundation* Distinguished Postdoctoral fellowship (DW), a *Carlsberg Foundation* instrument grant (MK), and a *Sapere Aude* advanced grant from the Danish Council for Independent Research | Natural Sciences (MK).

## Acknowledgments

We acknowledge Sofie Lindegaard Jakobsen, Erik Trampe, and the staff at Heron Island Research station for help and technical assistance during field work. Ingrun Schönberg and Jörg Wollenzin (Thorlabs GmbH, Lübeck, Germany) are thanked for technical advice on OCT system operation.

